# Development and evaluation of a dual target glycoconjugate vaccine against *Shigella sonnei*

**DOI:** 10.64898/2026.03.26.714513

**Authors:** Catherine L Hall, Tom L Flood, Simon Clare, Katherine Harcourt, Emily Kay, Stephen Baker, Brendan W Wren

## Abstract

**Background:** Shigellosis morbidity and mortality, combined with the increase in multidrug-resistant infections make *Shigella* vaccine development a global imperative. Glycoconjugate vaccines that couple immunogenic O-antigen to protein derived from *Shigella* may provide broader protection across *Shigella* species and serogroups. Such an approach also circumvents immunotolerance arising from repeated use of the same carrier. Here we use bioconjugation, exploiting an oligosaccharyltransferase (OST) enzyme to couple O-antigen and carrier protein *in vivo*, to generate a “double-hit” *Shigella* glycoconjugate vaccine.

**Method:** Glycoconjugates were synthesised in *E. coli* SDB1 cells expressing *S. sonnei* O-antigen, the OST PglS, and one of two *Shigella* carrier proteins. Recombinant glycoconjugate was purified using anion exchange chromatography and then used to immunise mice. Antibody responses were measured and compared by ELISA.

**Results:** When co-produced in *E. coli*, PglS was able to transfer the cloned *S. sonnei* O-antigen onto three carrier proteins, modified to accept glycans from the PglS transferase enzymes- the standard bioconjugate carrier ExoA and two immunogenic *Shigella*-specific outer membrane proteins, EmrK and MdtA. Production of MdtA or ExoA glycoconjugates for immunisation studies utilised successive rounds of anion exchange chromatography, to remove unglycosylated material and obtain highly purified glycoconjugate proteins for us in vaccination. Analysis of murine sera following immunisation revealed an IgG response was raised against both carrier protein and the *S. sonnei* O-antigen for each glycoconjugate.

**Conclusion:** A novel, conserved *Shigella* protein can be utilised as an effective carrier for the generation of a “double-hit”, immunogenic *Shigella* glycoconjugate vaccine that elicits IgG responses to both carrier protein and *S. sonnei* O-antigen.

## Introduction

Glycoconjugate vaccines have been instrumental in preventing some of the most serious and debilitating childhood infections such as pneumonia and meningitis (1, 2). Bacterial polysaccharides are regarded as key vaccine antigens but when delivered alone do not engage suitable T-cell responses, resulting in short-lived immunity and poor protection in children (3).

Covalent linkage of polysaccharides to a protein carrier enables their presentation via major histocompatibility complex class II to T-cells leading to T-cell activation and the generation of a robust, long-lasting immune response (3).

All currently licensed glycoconjugate vaccines are manufactured via chemical conjugation; polysaccharide and protein components are produced and purified separately before being chemically covalently linked (4). For decades, this approach has enabled the production of life-saving vaccines but is a costly and complex process requiring handling of pathogenic bacteria and multiple rounds of synthesis, purification and quality control (4, 5). Alternative means of synthesis such as bioconjugation could address these limitations via an *in vivo* method of glycoconjugate production. Here, glycoconjugate vaccines are produced heterologously in genetically modified strains of bacteria, such as *E. coli*, using bacterial oligosaccharyltransferase (OST) enzymes to couple polysaccharide and protein (Figure 1) (6–8). The simplicity of production compared to chemical conjugation has the potential to produce vast protein/glycan combinations and substantially reduce vaccine manufacturing costs (5).

**Figure 1.**
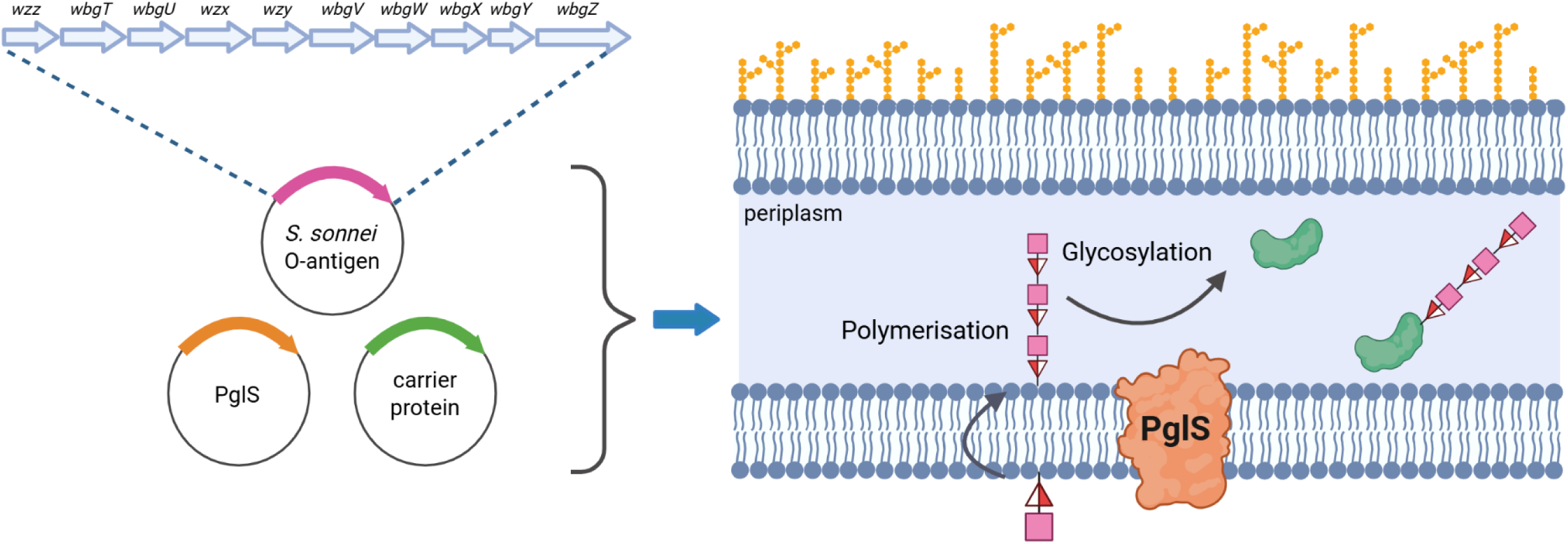
Schematic of bioconjugation for the production of a *Shigella sonnei*-specific glycoconjugate. Three plasmids are co-expressed in *E. coli* encoding a carrier protein, the oligosaccharyltransferase PglS and the *S. sonnei* O-antigen (SSOAg) (encoded by the *Plesiomonas shigelloides* O-antigen biosynthesis locus). The SSOAg (D-FucNAc4N-L-AltNAcA) is assembled on the cytoplasmic face of the inner membrane then flipped into the periplasm where it is polymerised then conjugated to a ComP-tagged carrier protein (green) by PglS (orange) to produce glycoconjugate. The red and white triangle and pink square are used here to represent D-FucNAc4N and L-AltNAcA, respectively. Schematic made using Biorender.

This is especially important considering many bacterial infections present the highest burden in low-and middle-income (LMIC) settings. Furthermore, a sub-set of bioconjugates have now advanced into clinical trials, such as S4V-EPA, a quadrivalent vaccine against *Shigella* infections (9).

Shigellosis is a leading cause of diarrhoeal disease, globally. Of the four *Shigella* species, *S. flexneri* typically causes the majority of infections in LMICs, particularly in children under five years (10). *S. sonnei* is more prominent in high income countries, although increasing global incidence driven by the spread of multidrug-resistant strains has led to its dominance in many regions (11). Shigellosis can range from self-limiting diarrhoea to acute bloody diarrhoea with fever and is estimated to cause 68,000 deaths in children under five, annually. For those who survive, many experience life-long consequences of infection (12, 13).

The increase in antimicrobial resistance alongside the morbidity and mortality of shigellosis makes *Shigella* a priority pathogen for vaccine development but despite numerous approaches, there remains no licensed candidate. This has in part been impeded by the lack of established correlates of protection, to guide the design of an efficacious vaccine. It is widely viewed that serum IgG against the *Shigella* lipopolysaccharide (LPS) is associated with protection from infection and as such, many vaccine candidates in clinical trials focus on targeting this antigen (14). These candidates include two chemically synthesised glycoconjugates, *S. flexneri* 2a and/or *S. sonnei* LPS conjugated to tetanus toxoid (15, 16) and S4V-EPA, a quadrivalent bioconjugate vaccine targeting *S. sonnei* and *S. flexneri* serotypes 2a, 3a or 6, with LPS from each conjugated to *Pseudomonas aeruginosa* exotoxin A (ExoA) (9). Alternative vaccine designs based on the *Shigella* LPS include Invaplex, a high molecular mass complex of Invasion protein antigens (Ipa) IpaB and IpaC, with *Shigella* LPS (17, 18) and alt-Sonflex1-2-3, a generalised module for membrane antigen (GMMA) based vaccine (19). This candidate uses a combination of outer membrane vesicles isolated from *S. sonnei* and three *S. flexneri* serotypes (1b, 2a and 3a) (20).

Notably, all these vaccine candidates incorporate LPS from multiple *Shigella* serogroups, which is necessitated by the structural variability of the *Shigella* O-antigens (outermost component of the LPS). *S. sonnei* harbours a single, conserved O-antigen structure, while there are multiple serotypes and sub-serotypes of *S. flexneri* (21). A vaccine targeting *S. sonnei* plus *S. flexneri* serotypes 2a, 3a and 6 is estimated to protect against approximately 80% of *Shigella* infections, based on the prevalence of the different serotypes as well as cross-reactivity between some serotypes of *S. flexneri* (12).

The use of multiple glycan antigens to achieve adequate serotype coverage of a vaccine has been demonstrated for licensed glycoconjugates against *Streptococcus pneumoniae* and *Neisseria meningitidis,* but does present a challenge to vaccine production (22). This could be overcome through use of the carrier protein not only for glycan presentation but as a serotype-independent immunogen, offering a means of expanding vaccine valency and dual stimulation of the immune response (23). However, to date, most glycoconjugate vaccines, both chemically and biologically synthesised, have relied on a small pool of well-characterised carrier proteins, such as detoxified tetanus and diphtheria toxoids for chemical conjugates and ExoA for S4V-EPA (6, 9). Repeated use of the same carrier proteins within glycoconjugate vaccines risks the development of carrier-induced epitope suppression and blunting of the immune response due to over exposure to the same protein antigens (23, 24). Design of such “double-hit” glycoconjugates requires the careful balancing of each antigen, to ensure the protein does not detract from the anti-polysaccharide antibody response and in turn, the polysaccharide does not mask important protein epitopes (1). Bioconjugation complements the design and development of such novel carrier proteins as the site of glycosylation can be precisely determined through the positioning of the OST recognition sequon. Although not yet widely explored, this “double-hit” approach has already been investigated for bioconjugate vaccines against *Streptococcus pneumoniae* and Group A *Streptococcus* (25, 26).

To date, *Shigella*-specific protein vaccine candidates have primarily focused on the immunogenic, well-characterised and conserved Ipa proteins, which are part of the type III secretion system (27). Immunisation of mice with *S. flexneri* 2a O-antigen conjugated to IpaB resulted in protection from lethal *S. flexneri* and *S. sonnei* challenge (28). Similar results have been obtained following immunisation with Invaplex, which contains both IpaB and IpaC. This demonstrates the potential of *Shigella-*specific proteins to provide heterologous protection from infection (17, 18).

Protein arrays have been widely utilised in reverse vaccinology as a high throughput means of identifying novel vaccine antigens for further characterisation and evaluation. de Alwis *et al.* used such an approach to identify a number of novel *Shigella* immuno-antigens via probing with acute and convalescent sera from patients with natural *Shigella* infection (29). Applying this understanding to the design of novel carrier proteins could facilitate the development of next-generation bioconjugate vaccines. In this study, we cloned the complete *S. sonnei* O-antigen pathway in *E. coli*, and used two immunogenic *Shigella* proteins, EmrK and MdtA as novel carriers for *Shigella* glycoconjugates. Both proteins, alongside the established bioconjugate carrier ExoA, could be conjugated to the *S. sonnei* O-antigen using the OST PglS. Evaluation of *S. sonnei-*specific glycoconjugates containing either MdtA or ExoA found both conjugates elicited a specific anti-protein and anti-O-antigen IgG response in a mouse infection model.

## 2. Methods

### 2.1 Bacterial strains and plasmids

A list of bacterial strains used in this study can be found in Table S1. *Escherichia coli* was routinely cultured using Luria-Bertani (LB) (Miller) agar or broth at 30°C, with shaking at 180 rpm for liquid cultures. Both LB and Terrific broth were used for large scale glycoconjugate production, as detailed in 2.6. Where necessary, media was supplemented with the following antibiotics for plasmid maintenance; ampicillin (100 µg ml^-1^), tetracycline (20 µg ml^-1^), chloramphenicol (30 µg ml^-1^) and kanamycin (50 µg ml^-1^).

### 2.2 Construction of the Plesiomonas shigelloides O17 O-antigen expression plasmid

A list of plasmids and oligonucleotides used in this study can be found in Tables S2 and S3, respectively. The sequence of all plasmids was confirmed using long-read sequencing.

Cloning and synthesis of the *S. sonnei* O-antigen (SSOAg) used gDNA from *Plesiomonas shigelloides* O17 (NCTC10360), as the two harbour identical O-antigen structures (30). A 12.2 Kb region of the *P. shigelloides* O17 O-antigen, from *wzz* to *wbgZ* was amplified using Q5 polymerase with primers PsOFApal and PsORApal. Primers included ApaI restriction sites to enable insertion of the O-antigen fragment into pBBR1MCS-3 using restriction ligation cloning.

### 2.3 Construction of carrier protein and PglS expression plasmids

Carrier proteins used for glycoconjugate production were comprised of an N-terminal DsbA sequence for trafficking to the periplasm, protein coding sequence, C-terminal ComP tag (amino acids 29 to 145) for glycosylation with the PglS OST (31) and a final C-terminal 6XHistag for protein purification (Figure S1). The detoxified, C-terminal fragment of ExoA used in this study has been described elsewhere (32, 33). The sequence of ExoA used to construct pEC415_ExoAComP also contains two inserted glycosylation sequons recognised by the PglB OST (32). These were inactivated in this study by replacing the glycosylated asparagine residues with glutamine (34). The amino acid sequences of EmrK and MdtA were derived from *S. sonnei* and modified to remove the N-terminal native signal peptide, resulting in expression of amino acids 38 to 387 for EmrK and 66 to 453 for MdtA.

Carrier proteins were initially designed with two ComP tags, flanking the protein coding sequence at both the N- and C-termini (Figure S1). These dual tagged carrier proteins were ordered as gBlocks, with additional 5’ and 3’ overhangs for insertion via Gibson assembly into pEC415, amplified using primers pEC415DsbA_f and r. gBlocks did not include the DsbA signal sequence as this was already present within the pEC415 plasmid backbone, located 9 base pairs downstream of the ribosome binding site. Due to absent or very low protein expression from these constructs (data not shown), the N-terminal ComP tag was removed by inverse PCR then plasmids ligated using the KLD Enzyme Mix (NEB). Primers for inverse PCR were as follows; ExoA_NComPdel_f and r for pEC415_ExoAComP, EmrK_NComPdel_f and r for pEC415_EmrKComP and MdtA_NComPdel_f and r for pEC415_MdtAComP. ExoA and MdtA with no ComP tag(s) were ordered as gBlocks and inserted using Gibson assembly into pEXT20, amplified using primers pEXT20_noComP_f and r (Figure S1).

To generate pEXT22_pglS, the nucleotide sequence of *Acinetobacter baylyi pglS* was ordered as a gBlock. A minor number of base changes were required to permit gBlock synthesis, but these did not alter the amino acid sequence. EcoRI and BamHI restriction sites were included at the 5’ and 3’ of the sequence, respectively, for insertion into pEXT22 using restriction ligation cloning.

### 2.4 Chromosomal deletion of mdtA from E. coli W3110

*mdtA* was deleted from the *E. coli* W3110 chromosome using lambda red recombineering (35). The deletion cassette was comprised of two 120 bp homology arms, representing the DNA sequences immediately upstream and downstream of *mdtA.* Within the homology arms was a chloramphenicol cassette flanked by two *dif* recombinase sites (36). This construct was ordered as a gBlock and inserted using Gibson assembly into pKD4, amplified using primers pKD4MdtA_f and r. Gibson reactions were dialysed into dH_2_O using 0.45 µM membrane filters before transformation into electrocompetent TransforMax *E. coli* EC100D (*pir* +) (Biosearch Technologies). Transformants were selected on LB agar with chloramphenicol and ampicillin and extracted plasmids confirmed with long-read sequencing, to produce pKD4_mdtA.

pKD4_mdtA was used as the template for amplification of the deletion cassette, from the 5’ of homology arm one to the 3’ of homology arm two, using primers MdtAcassette_f and r. Purified amplicon was used to transform electrocompetent *E. coli* W3110 harbouring the lambda red recombineering plasmid, pKD46. Transformants were selected on LB agar with chloramphenicol and colonies screened for cassette integration by PCR, using primers MdtAscreen_f and r.

PCR-confirmed, chloramphenicol-resistant integrants were passaged on LB agar only at 37°C, to facilitate loss of the temperature sensitive pKD46 and for removal of the chloramphenicol cassette, using the endogenous *E. coli* recombination system, Xer, which recognises the *dif* recombinase sites (36). Chloramphenicol and ampicillin sensitive colonies were screened for loss of the chloramphenicol cassette by PCR, using the same screening primers as above, and confirmed using long-read sequencing.

### 2.5 Small scale glycoconjugate production and purification

To screen for SSOAg production, strains were grown overnight in 5 ml LB, shaking at 37°C then harvested by centrifugation at 17,000 xg for 10 mins at 4°C. To screen for glycoconjugate production, a starter culture was grown for 16-18 hours and used to inoculate 40 ml LB at a 1:100 dilution, then incubated shaking at 30°C. When OD_595_ reached 0.5 to 0.7, 0.5 mM IPTG was added to induce OST expression. Following overnight incubation, cultures were induced with 0.2% L-arabinose for carrier protein expression then incubated at 25°C for a further 8 hours. Cells were harvested by centrifugation at 17,000 xg for 10 mins at 4°C.

For protein extraction and purification, cell pellets were resuspended in 2 ml binding buffer (50 mM Tris-HCl, 300 mM NaCl, pH 8.0) and added to lysing matrix B for lysis on a FastPrep homogeniser. Lysates were centrifuged twice at 17,000 xg for 5 mins to remove cell debris then the supernatant was incubated with 2% Triton X-114 at 4°C, rotating, for 1 hour or overnight.

Triton X-114 was removed by heating at 37°C for 10 mins then centrifugation at 17,000 xg for 10 mins. The upper phase was isolated and 6XHis-tagged proteins purified using nickel affinity chromatography and stored in elution buffer (50 mM Tris-HCl, 300 mM NaCl, 300 mM imidazole, pH 8.0).

### 2.6 Glycoconjugate production and purification for immunogenicity studies

Glycoconjugate and unglycosylated protein to be used in immunogenicity studies were produced in *E. coli* strain SDB1. SSOAg-ExoAComP glycoconjugate was produced using pBPSO (SSOAg), pEXT22_pglS (PglS) and pCH4 (ExoAComP) (31). The SSOAg-MdtAComP glycoconjugate was produced using pBPSO, pACT3_pglS (PglS) and pEC415_MdtAComP (MdtAComP). Unglycosylated protein was produced using pCH4, pEC415_MdtAComP (proteins with ComP), pEXT20_ExoA or pEXT20_MdtA, only (proteins without ComP).

#### 2.6.1 Bacterial cell growth

All cultures were grown in batches of 2 L media in a 5 L shake flask. For production of SSOAg-ExoAComP glycoconjugate and all unglycosylated proteins, LB starter cultures were grown at 30°C for 16-18 hours and used to inoculate 2 L LB at 1:100. Cultures were incubated at 30°C, 150 rpm and induced with 0.5 mM IPTG at OD_595_ 0.5-0.7, then incubated overnight. For the SSOAg-MdtAComP glycoconjugate, starter cultures in Terrific broth (Table S4) were diluted 1:50 into 2 L Terrific broth. At OD_595_ 0.6-0.8, cultures were induced with 0.5 mM IPTG and temperature reduced from 30°C to 25°C. After two hours, 0.2% L-arabinose was added then cultures incubated overnight. Cells were harvested at 7500 xg for 30 minutes at 4°C and cell pellets frozen at -20°C.

#### 2.6.2 Protein Extraction

For periplasmic extraction of SSOAg-ExoAComP, each 2 L cell pellet was resuspended in approximately 160 ml periplasmic extraction buffer (30 mM Tris-HCl, 20% sucrose, 1 mM EDTA, 1 mg/ml lysozyme, pH 8.0) and incubated at 4°C for 1 hour on a rolling platform. Lysates were incubated for a further 30 mins at room temperature with 10 units/ml of Benzonase nuclease and 2 mM MgCl_2_ then centrifuged for 1 hour, 31,000 xg, 4°C. Supernatant was passed through a 0.2 µM filter then dialysed into binding buffer overnight at 4°C.

SSOAg-MdtAComP glycoconjugate and all unglycosylated proteins were extracted using whole cell lysis. Cell pellets from 2 L cultures were resuspended in 100-200 ml binding buffer and lysed using a pre-chilled pressure cell homogeniser. Lysates were then incubated at room temperature for 30 mins with 1 mg/ml lysozyme and 10 units/ml of Benzonase nuclease then centrifuged for 1 hour, 31,000 xg, 4°C.

To remove lipid and lipid-linked glycan from the cell extracts, 2% Triton X-114 was added to chilled, filtered lysates and incubated at 4°C for 1 hour or overnight on a rolling platform. Triton X-114 was removed by incubating at 37°C for 10 mins then centrifugation at 3900 xg. The upper phase was isolated and centrifuged again at 3900 xg for 5 mins then upper phase saved for protein purification.

#### 2.6.3 Ni-NTA affinity chromatography

Nickel affinity chromatography was performed using Ni-NTA resin in 10 ml polypropylene columns. Resin was prepared by washing with 10 column volumes (CV) of dH_2_O, 5 CV of elution buffer and 10 CV of binding buffer. Prepared resin was combined with Triton X-114-treated lysates and incubated at 4°C for 1-2 hours on a rolling platform before loading onto the column. The flowthrough fraction was captured and reserved then resin washed three times with 10 CV wash buffer (50 mM Tris-HCl, 300 mM NaCl, 10 mM imidazole, pH 8.0). Resin was incubated for 2 mins with 1 CV elution buffer before elution, this was repeated a total of 5 times. For the SSOAg-ExoAComP glycoconjugate and ExoA and MdtA only, the reserved flowthrough fraction was passed over the re-equilibrated resin again and method repeated one to two more times.

Nickel affinity-purified samples were stored either at 4°C or with 20% glycerol at -80°C.

#### 2.6.4 Anion Exchange Chromatography

Nickel affinity purified samples were buffer exchanged into AEX start buffer using PD-10 desalting columns (Cytiva) or overnight dialysis. For the AEX start buffers, BisTris pH 6.0 was used for purification of SSOAg-ExoAComP glycoconjugate and unglycosylated ExoAComP or ExoA and Tris pH 8.6 for SSOAg-MdtAComP glycoconjugate and unglycosylated MdtAComP or MdtA. AEX was performed on an AKTA Go Protein Purification System (Cytiva), using a 1 ml Resource Q column. Glycoconjugate and unglycosylated protein was eluted from the column using AEX start buffer plus a NaCl gradient up to 500 mM. Fractions of interest, as confirmed by SDS-PAGE, Imperial stain and Western Blotting, were pooled and transferred back into AEX start buffer using overnight dialysis, PD-10 desalting columns or dilution into start buffer. AEX was repeated 3-4 times for glycoconjugate purification and 1-3 times for protein only. Final fractions not undergoing further purification were transferred into endotoxin-free, sterile phosphate buffered saline, pH 7.4, using PD-10 columns, quantified using a Qubit 3 and lyophilised.

### 2.7 SDS-PAGE and Western Blotting

Glycoconjugate and protein samples were mixed with 1 x LDS loading buffer (Thermo Fisher Scientific) and 1-10 mM DTT then heated for 10 mins at 70°C or 5 mins at 95°C. 20-40 µl was loaded onto a 4-12% BisTris gel and run in MOPs buffer. For glycan only samples, 5 ml cell pellets were resuspended in PBS to OD 1.5 then 500 µl removed and boiled for 10 mins before mixing with LDS loading buffer as above. Gels were transferred to nitrocellulose membrane using an iBlot3 dry blotting system. Following transfer, SDS-PAGE gels were washed with dH_2_O for 10 mins, incubated with Imperial stain for 1 hour then destained using multiple washes in dH_2_O. Imperial-stained gels were imaged on a Licor Odyssey M. The same method for was used for Imperial staining of those SDS-PAGE gels not also being used for Western Blotting.

Membranes were washed 3 x 5 mins with PBS Tween 0.1% then incubated for 45 mins to 1 hour with primary antibodies- Mouse anti-*S. sonnei* monoclonal antibody (Abnova) at 1:2000 and Rabbit anti- 6xHistag polyclonal antibody (Invitrogen), at 1:5000 dilution. Membranes were washed as described then incubated for 30-45 mins with secondary antibodies- IRDye680 Goat anti-mouse IgG and IRDye800CW Goat anti-rabbit IgG (LI-COR). Membranes were washed for a final time before imaging using the LI-COR Odyssey M.

For densitometry analysis of glycoconjugate production, Western blot images were converted to grey scale and exported into software (v1.54g). A defined rectangle of the same size was placed around the glycoconjugate band(s) for each protein and mean grey value measured. An empty control lane was used to determine the background, and this was subtracted from each glycoconjugate reading. Statistical significance was determined from three biological replicates using Welch’s ANOVA with Games-Howells’ multiple comparison test.

### 2.6 Animal study

All animal experiments were performed in accordance with UK Home Office regulations under project licenses PP2473276 following approval from the local University of Cambridge Animal Welfare and Ethical Review Body.

Six-week old, C57BL6J female mice were purchased (Charles River). Mice were housed in GM500 Individually Ventilated Cages- Biocontainment Systems (Tecniplast) and maintained on individual air handling units kept at between 19 to 23°C and 45–65% humidity; each cage received 60 air volume changes per hour. Mice were given access to food and water ad libitum and maintained on a 12-hour light/dark cycle. Aspen wood chip was used as the base bedding with nestlets and “fun tunnels” for environmental enrichment (Datesand). Mice were housed in groups of no more than five adults per cage and acclimatised for one week prior to immunisation. Welfare assessments were carried out daily as part of the morning check. This involved moving the cage forward on the rack runners and observing the mice in their micro-environment. Any abnormal signs of behaviour or physical clinical signs of concern were reported. “In hand” weekly health assessments were also performed, and every cage was removed from the rack and each mouse given a full “head to toe” health check. All persons performing welfare checks on animals were trained and assessed as competent by qualified individuals.

Mice (n= 7 per group) were immunised sub-cutaneously in the flank with either glycoconjugates (SSOAg-ExoAComP or SSOAg-MdtAComP), carrier protein only (ExoAComP or MdtAComP) or PBS only. All glycoconjugates and carrier proteins were tested individually, not in combination. Mice were immunised with 10 µg of protein per dose in PBS and adjuvant Alhydrogel (1:1) (InvivoGen) on days 0, 14 and 28. On day 35 mice were killed by terminal anaesthesia followed by exsanguination and cervical dislocation. Blood was collected into sterile 1.5 ml microcentrifuge tubes and left to clot for 30 mins. The serum was removed by centrifugation at 21,000 x g for 5 mins and collected into a fresh tube and stored at -20°C until used in assays.

### 2.7 Anti-protein ELISAs

ELISAs were used to measure murine IgG responses to ExoA and MdtA . 96-well Maxisorp plates were coated with 0.2 µg/well of ExoA or MdtA, or PBS only. Following overnight, static, incubation at 4°C, plates were washed four times with PBS Tween 0.05% then 100 µl/well of blocking buffer (PBS Tween 0.05%, 5% skimmed milk) was added and plates incubated at 500 rpm, room temperature for 1 hour. Plates were washed three times with PBS Tween 0.05% then 100 µl/well sera was added and plates incubated at 500 rpm, room temperature for 1 hour. Sera from each mouse was tested individually in technical duplicate and serially diluted in PBS Tween 0.05%, using a 6-point, 3-fold dilution series. Plates were washed four times with PBS Tween 0.05% then incubated with 100 µl/well of HRP-conjugated Goat anti-mouse IgG, at 1:10,000 dilution, room temperature for 1 hour. Plates were washed four times with PBS Tween 0.05% then incubated with 100 µl/well TMB substrate. The reaction was stopped after 8 mins using 100 µl/well of 1 M sulphuric acid then OD_450_ and OD_570_ read using a plate reader.

Background subtractions were made using the OD_570_ and no sera control readings. A coefficient of variation (CV) of <20% was considered acceptable for inter-plate and intra-plate variability. Inter-plate variability was measured using a pool of sera from mice immunised with ExoAComP or MdtAComP only, tested against their respective antigen on each plate and intra-plate variability determined using duplicate testing of individual sera samples. OD_450_ values obtained from serial dilution of each sera sample was plotted on a 4-parameter logistic standard curve in GraphPad Prism and used to determine the reciprocal of the sera dilution that gave an OD 0.5. The reciprocal determined for mice immunised with PBS only was greater than the lowest dilution of serum tested (1/600) therefore a serum dilution just outside of this range (599) was used for visualisation and data analysis. Statistically significant differences in IgG responses against each protein was determined by Kruskal-Wallis test with Dunn’s multiple comparison test.

For the anti-protein ELISAs measuring the IgG response to the carrier proteins with and without ComP, the above protocol was followed with minor amendments. Plates were coated with 0.5 µg/well of ExoAComP, ExoA, MdtAComP or MdtA, or PBS only and probed with pooled sera from mice immunised with either ExoAComP or MdtAComP only, at a 1 in 100,000 dilution. Sera was tested in technical triplicate and OD_450_ reported after the background subtracted using values from a no antigen control.

### 2.8 Anti-glycan ELISAs

Whole cell ELISAs were used to measure murine IgG responses to the SSOAg. A starter culture of *E. coli* W3110Δ*mdtA* harbouring either pBPSO, encoding the SSOAg locus or empty vector (pBBR1MCS-3) were grown for 16-18 hours and used to inoculate 20 ml LB at a 1:100 dilution. Cultures were incubated shaking at 37 °C until OD_595_ reached 0.6 then centrifuged at 4300 xg for 10 mins. Cells pellets were washed three times in 20 ml Carbonate buffer (15 mM Na_2_CO_3_ and 35 mM NaHCO_3_), OD_595_ adjusted to 0.6 then diluted one in ten in Carbonate buffer and used to coat 96-well Maxisorp plates, 100 µl/well, which equated to approximately 3-4 x 10^6^ cells per well. Coated plates were air-dried in a microbiological safety cabinet then fixed with 100 µl/well methanol and plates air-dried again then stored for up to 1 week.

For the anti-glycan ELISA, coated plates were washed four times with PBS Tween 0.05% then 100 µl/well of blocking buffer (PBS Tween 0.05%, 10% skimmed milk) was added and plates incubated at 500 rpm, room temperature for 2 hours. Plates were washed three times with PBS Tween 0.05% then 100 µl/well sera were added and plates incubated at 500 rpm, room temperature for 2 hours. Pooled sera from each group were serially diluted in PBS Tween 0.05% + 1% skimmed milk, using a 6-point, 3-fold dilution series. Plates were washed four times with PBS Tween 0.05% then incubated with 100 µl/well of HRP-conjugated Goat anti-mouse IgG, at 1:10,000 dilution, room temperature for 1 hour. Plates were washed four times with PBS Tween 0.05% then incubated with 100 µl/well TMB substrate. The reaction was stopped after 13 mins using 100 µl/well of 1 M sulphuric acid then OD_450_ and OD_570_ read using a plate reader.

Pooled sera was tested in technical triplicate and background subtractions were made using the OD_570_ and no sera control readings. A pool of sera from mice immunised with SSOAg-ExoAComP was tested against wells coated with W3110Δ*mdtA* + pPBSO on each plate, to measure inter-plate variability, for which a CV of <20% was considered acceptable. A CV of <20% was also deemed acceptable for intra-plate variability, measured using triplicate testing of pooled sera samples. OD_450_ values obtained from serial dilution of each serum sample was plotted on a 4-parameter logistic standard curve in GraphPad Prism and used to determine the reciprocal of the serum dilution that gave an OD 0.2.

## 3. Results

### 3.1 Generation of Shigella-specific bioconjugates using PglS

The *S. sonnei* O-antigen (SSOAg) constitutes a two component repeat unit of a reducing end 2-acetamido-4-amino-2, 4-dideoxy-D-fucose (D-FucNAc4N) linked to 2-amino-2-deoxy-L-altruronic acid (L-AltNAcA) (21, 37). This structure is distinct from those found in other *Shigella* and *E. coli* species and was acquired via horizontal gene transfer from *P. shigelloides* O17, which harbours an identical O-antigen (30). Using the nucleotide sequence from the *P. shigelloides* O-antigen biosynthesis locus (pBPSO), we recombinantly produced the SSOAg in *E. coli* SDB1, as confirmed by Western blot using a *S. sonnei-*specific monoclonal antibody (Figure 2A).

**Figure 2.**
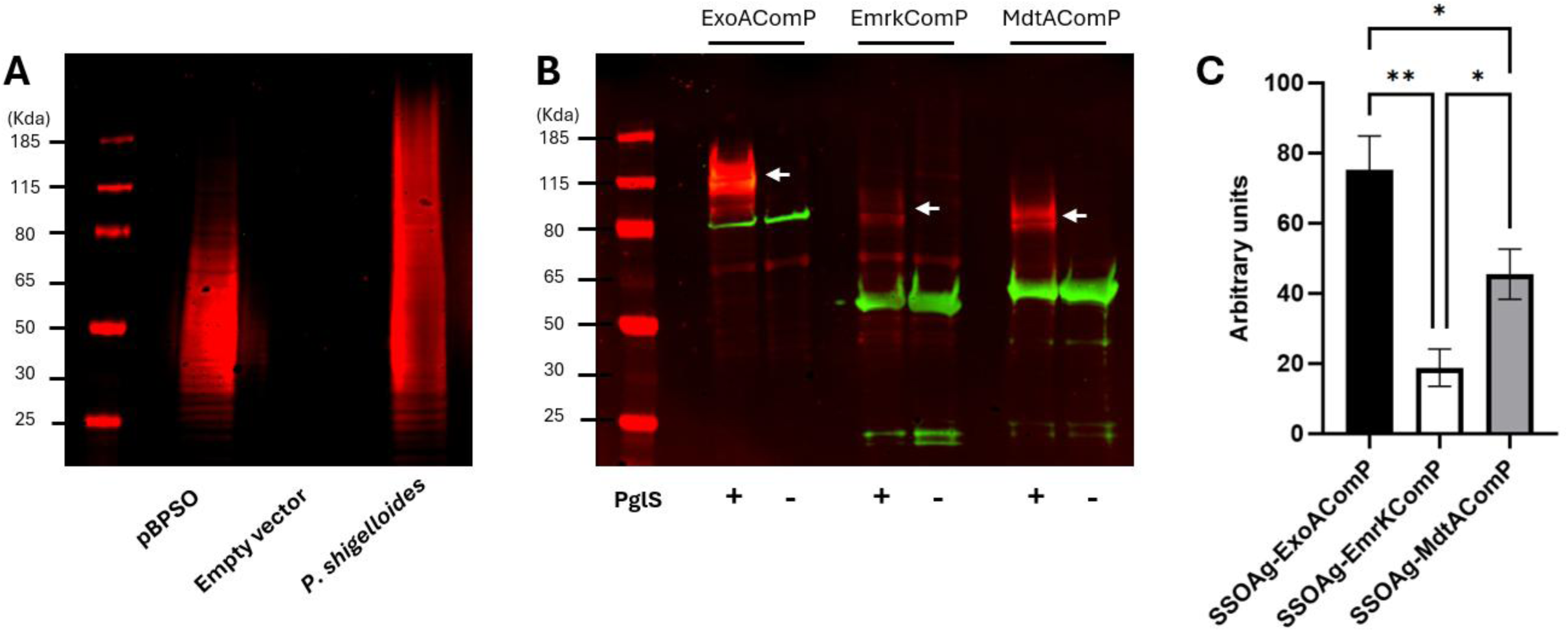
Glycosylation of *Shigella*-specific carrier proteins using the oligosaccharyltransferase (OST) PglS. (A) Recombinant production of the *Shigella sonnei* O-antigen (SSOAg) in *E. coli* using the O-antigen biosynthesis locus from *Plesiomonas shigelloides* (pBPSO). Cell lysates from either *E. coli* carrying pBPSO or empty vector (pBBR1MCS-3) or *P. shigelloides* alone, were resolved by SDS-PAGE and detected via Western blot using anti- SSOAg antibody (red). (B) ExoA, EmrK and MdtA were modified with a C-terminal ComP tag for glycosylation by the PglS OST and co-expressed in *E. coli* SDB1 with the *S. sonnei* O-antigen and PglS, or a no-OST control (pSta313). Glycosylated and unglycosylated carrier proteins were resolved by SDS-PAGE and detected via Western blot using anti-6xHistag antibody (green) and SSOAg antibody (red). (C) Densitometry analysis of *S. sonnei* glycoconjugate production for the three carrier proteins. ImageJ software was used to quantify glycoconjugate production. Statistical significance was determined from three biological replicates using Welch’s ANOVA with Dunnett’s T3 multiple comparisons test, \*\**p=*<0.01, \**p=*<0.05. SSOAg, *S. sonnei* O-antigen.

OSTs can exhibit substrate specificity in the glycans they transfer and the SSOAg harbours an atypical reducing end D-FucNAc4N, so we attempted to conjugate the glycan using OSTs from N-linked and O-linked glycosylation systems, namely *C. jejuni* PglB and *Acinetobacter baylyi* PglS, respectively (Figure 1) (8, 31, 33, 38). Western blot analysis using an anti-SSOAg antibody, revealed glycoconjugate production in strains producing the SSOAg with PglS and its carrier protein ExoAComP, with no glycoconjugate production observed using PglB and its compatible carrier, ExoA (Figure S2). Glycosylated ExoAComP migrated between 110 and 120 KDa, compared to the unglycosylated ExoAComP at approximately 80 KDa. Glycosylated ExoAComP presented as a less defined band compared to the unglycosylated protein, possibly representing different sizes of glycoconjugate. As this carrier protein only harbours a single glycosylation site, this is likely a result of varying polymer lengths of the attached SSOAg. These results confirmed PglS as a viable OST for synthesis of *S. sonnei*- specific glycoconjugates.

We next investigated whether immunogenic *Shigella*-specific proteins could be used as novel glycoconjugate carrier proteins. Those selected, EmrK and MdtA, are both components of efflux pumps and conserved across *S. sonnei* and *S. flexneri* (29). Both were sero-reactive in a *Shigella*-specific protein array, with a significant increase in IgM, IgG and IgA responses to both proteins when comparing acute and early convalescent sera from patients with confirmed *Shigella* diarrhoea (29). Both proteins were modified with a C-terminal ComP tag (Figure S1), and when co-expressed with the SSOAg and PglS, produced *S. sonnei*-specific glycoconjugate (Figure 2B). Densitometry analysis of glycoconjugate production (Figures S3) revealed that the highest glycoconjugate yield was obtained using ExoAComP as the carrier protein. Both ExoAComP and MdtAComP produced significantly more glycoconjugate than EmrKComP, based on total yield from a 40 ml culture (Figure 2C). Therefore, only ExoAComP and MdtAComP were taken forward for immunogenicity studies.

### 3.2 Purification of glycoconjugates for immunogenicity studies

SSOAg glycoconjugates were purified for immunogenicity studies using a combination of Ni-NTA and anion exchange chromatography (AEX). Although an initial Ni-NTA purification step was sufficient to capture *S. sonnei* glyconjugate from the cell lysate, some contaminating proteins remained, in addition to a large proportion of unglycosylated protein. As glycosylation can alter the isoelectric point of a protein (39), AEX was used to further purify the glycoconjugates as this separates macromolecules based on charge.

Buffers with a pH higher than the predicted pI of each protein were used to confer a negative charge, and enable protein binding to the positively charged AEX resin. Using Tris buffer pH 8.6, unglycosylated MdtAComP eluted from the column before the glycoconjugate but this did not completely separate the two species, therefore sequential rounds of AEX were employed to remove unglycosylated protein and yield pure glycoconjugate (Figure 3). For the production of SSOAg-ExoAComP, multiple rounds of AEX in BisTris buffer pH 6.0 were used and the glycoconjugate eluted from the column before the unglycosylated material (Figure S4).

**Figure 3.**
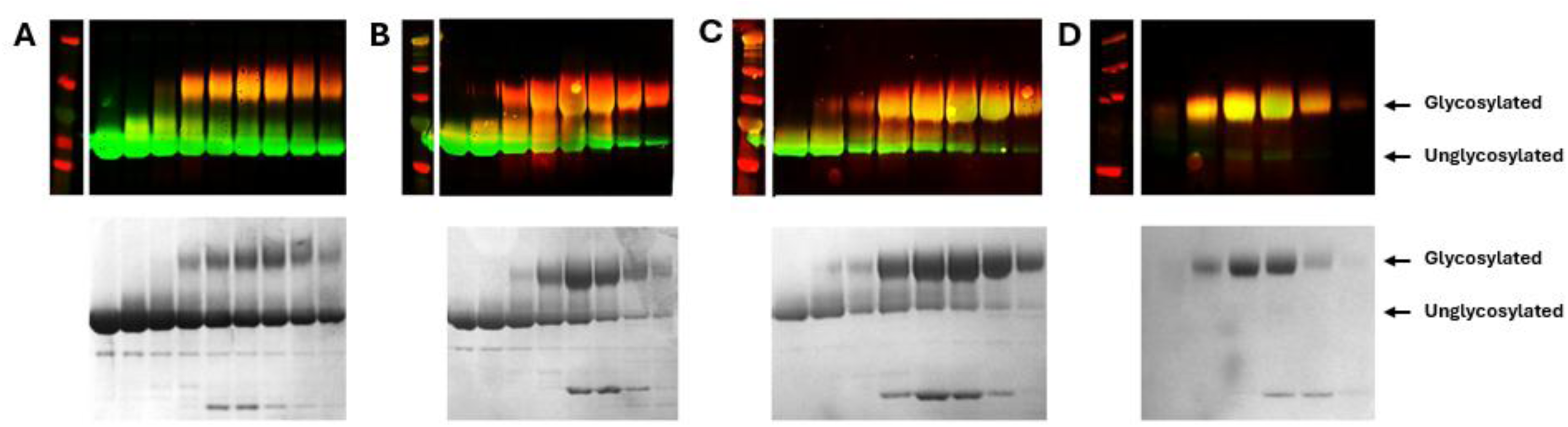
Sequential anion exchange chromatography (AEX) for purification of *S. sonnei-*specific glycoconjugates. Nickel-affinity purified MdtAComP conjugated to the *S. sonnei* O-antigen (SSOAg) was further purified via AEX for separation of glycosylated and unglycosylated protein using an increasing NaCl gradient. Elution fractions from each round of AEX were resolved by SDS-PAGE and visualised by Western blot (top panel) and Imperial staining (bottom panel). Fractions containing the most glycoconjugate were pooled and AEX repeated. Images A to D represent sequential rounds of AEX, from first to last, with each well containing a different elution fraction. Western blots were visualized using anti-6xHistag antibody (green) and anti-*S. sonnei* O-antigen (SSOAg) antibody (red/orange).

### 3.4 Anti-protein immune response

The immunogenicity of the SSOAg glycoconjugates was evaluated using C57BL/6j mice, immunised following a three-dose schedule of 10 µg protein per dose. Each glycosylated and unglycosylated protein was tested individually, alongside a PBS only control group. All were antigens were found to be well-tolerated.

Initial screening of the anti-carrier protein IgG response revealed all sera samples, except those from PBS-immunised mice, recognised both ExoAComP and MdtAComP. As both proteins share the ComP tag, it was hypothesised that the observed cross-reactivity was attributable to anti-ComP antibodies. Production of ExoA and MdtA only, without ComP (Figure S1B), eliminated this cross-reactivity and an anti-protein IgG response was only detected when probed with sera from mice immunised with the cognate carrier protein (Figure S5). As such, ExoA and MdtA only, without ComP tags, were used for characterisation of the anti-protein antibody response. Immunisation with both glycosylated and unglycosylated carrier proteins elicited a protein-specific IgG response. A significantly increased anti-ExoA IgG response was observed in mice vaccinated with either glycosylated or unglycosylated ExoAComP, compared to the PBS control (Figure 4, Figure S6). A comparable result was found for both MdtAComP vaccines against MdtA

**Figure 4.**
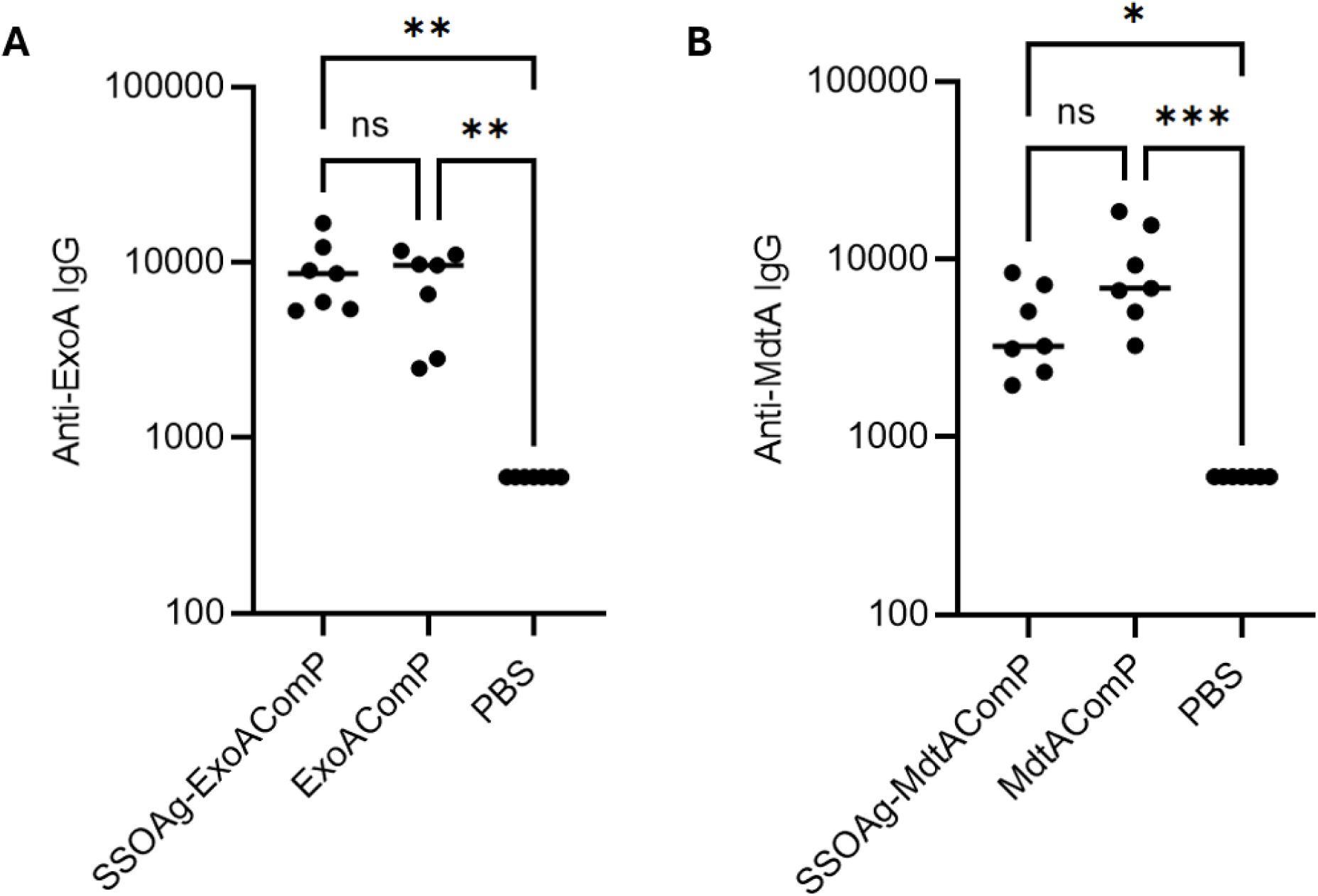
Anti-carrier protein antibody response to ExoA and MdtA. ELISA quantification of IgG response to ExoA and MdtA in mice immunised with glycosylated or unglycosylated ExoAComP or MdtAComP. A 6-point, 3-fold dilution series of individual serum samples from each group were tested in duplicate against purified ExoA or MdtA and IgG response reported as the reciprocal serum dilution that gave an OD 0.5. The reciprocal for PBS-immunised mice was outside of the tested dilution series (>1/600) so a value of 599 was used for visualisation and data analysis. Statistical significance was determined by Kruskal Wallis with Dunn’s multiple comparison test, \**p=*<0.05, \*\**p=*<0.01, \*\*\**p=*<0.001. SSOAg, *S. sonnei* O-antigen.

only. No significant difference in antibody response was observed between glycosylated versus unglycosylated protein, although the mean IgG response to unglycosylated MdtAComP was higher than that for the glycosylated protein. Overall, this data indicates glycosylation does not abolish the ability of either carrier to induce a protein-specific antibody response.

### 3.5 Anti-SSOAg immune response

To characterise the anti-glycan antibody response, serum samples were probed against whole *E. coli* cells producing the SSOAg. Given their similarity, both *Shigella* and *E. coli* encode a copy of the MdtA protein, which share a high degree of amino acid sequence homology. Although MdtA is a putative periplasmic protein, it may be accessible on the cell surface, therefore, to avoid interference from anti-MdtA antibodies, *mdtA* was deleted from the *E. coli* strain W3110, using lambda red recombineering (Figure S7). W3110Δ*mdtA* carrying either pBPSO (SSOAg) or pBBR1MCS-3 (empty vector) was then used to assess the anti-SSOAg IgG response. It was necessary to omit one serum sample from the PBS group as this was highly reactive against all *E. coli* cells, irrespective of glycan expression.

Serum IgG responses against the SSOAg were detected for both glycoconjugates (Figure 5). When tested against *E. coli* expressing the SSOAg, pooled sera from the SSOAg-ExoAComP immunisation group demonstrated an eight-fold increase in signal compared to ExoAComP only (Figure S8). Similarly, the SSOAg-MdtAComP signal was approximately two-fold higher compared to MdtAComP only. Furthermore, the anti-SSOAg response for glycosylated ExoAComP was approximately three-fold higher than that for glycosylated MdtAComP. Collectively, we have demonstrated the first use of PglS to generate novel, immunogenic *S. sonnei* glycoconjugates and use of *Shigella*-derived proteins as alternative, species-specific glycan carriers.

**Figure 5.**
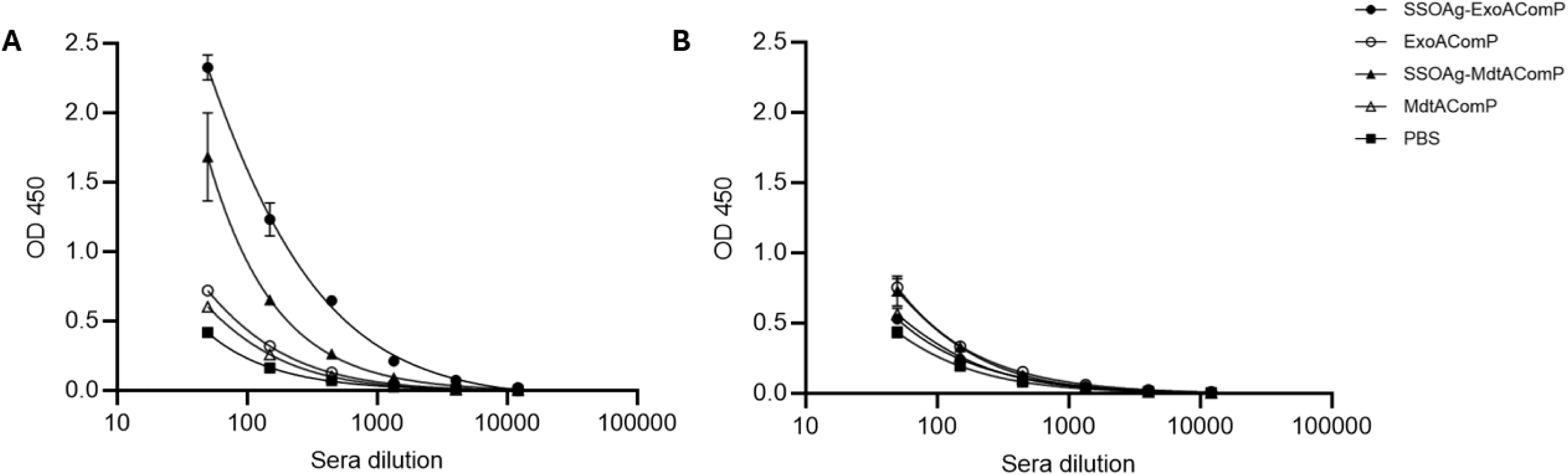
Antibody response to *S. sonnei* O-antigen. ELISA quantification of IgG response to whole *E. coli* cells producing (A) *S. sonnei* O-antigen or (B) no O-antigen, in mice immunised with glycosylated or unglycosylated ExoAComP or MdtAComP. OD_450_ is reported for the 6-point, 3-fold dilution series of pooled serum from each group and data fitted with a four-parameter logistic curve. Error bars represent standard deviation for three technical replicates. SSOAg, *S. sonnei* O-antigen.

## 4. Discussion

The application of bioconjugation for the synthesis of *Shigella* glycoconjugates offers flexibility in design and a means of low-cost vaccine manufacturing, a current imperative given the burden of shigellosis in LMICs. The identification of PglB and characterisation of its relaxed glycan substrate specificity paved the way for bioconjugate vaccine development (7, 8, 40). However, a reliance on wild-type PglB alone restricts antigen choice to only those glycans permissible for transfer by the OST. The identification of novel OSTs and structural-guided mutagenesis of existing ones has expanded the glycoengineering toolbox and consequently the repertoire of glycans available for bioconjugation (33, 41–43).

Here, we show biosynthesis of the *S. sonnei* O-antigen (SSOAg) in *E. coli* cells, by adopting the locus from *P. shigelloides* O17, which encodes an identical O-antigen (30). This is the first report of efficient, heterologous production of the SSOAg, which could be used for the development of both chemical and bioconjugates. It is also the first reported use of an *O*-linked oligosaccharyltransferase (OST) PglS, for a bioconjugate-derived *Shigella* vaccine. PglS was used to transfer SSOAg, which harbours an unusual reducing end monosaccharide, FucNac4N, to generate “double-hit” *S. sonnei-*specific glycoconjugates. Generation of this prototype vaccine was enabled by modification of *Shigella* proteins into permissible substrates for PglS-mediated glycosylation and as such novel glycoconjugate carriers. Immunisation with these glycoconjugates stimulated production of specific anti-protein and anti-glycan antibodies in a mouse infection model.

The majority of *S. sonnei* and/or *S. flexneri* O-antigen targeting glycoconjugate vaccines currently in clinical trials use established protein carriers such as tetanus toxoid or in the case of the S4V-EPA bioconjugate, ExoA (10, 16, 24, 44, 45). Species-specific carrier proteins could circumvent issues of immunotolerance from repeated exposure to the same carrier proteins, provide a means of dual immune stimulation and generation of serotype-independent immunity using a common antigen (6, 23, 46). The latter has already been demonstrated in clinical trials using Invaplex, a high molecular mass complex of two highly conserved *Shigella* proteins IpaB and IpaC and one or more purified *Shigella* LPS. In a pulmonary lung model, mice immunised with Invaplex containing IpaB, IpaC and purified *S. sonnei* LPS were protected from challenge with both *S. sonnei* and *S. flexneri* 2a, demonstrating the potential for protein-induced heterologous immunity (18). However, as with chemically conjugated glycoconjugate vaccines, these complexes require multiple expression and purification steps of each antigen to yield the final product.

This study is the first demonstration of the use of bioconjugation to generate a “double-hit”, shigellosis vaccine with the novel *Shigella*-specific carrier protein, MdtAComP. Both MdtA EmrK were previously identified as immunogenic in a protein array using early convalescent sera from patients with Shigellosis (29). Both could be glycosylated in this study when fused with a C-terminal ComP tag, the substrate of PglS (31). However, glycoconjugate yield varied between carrier proteins, despite their fusion to identical C-terminal tags. The structure of the carrier protein could potentially influence this, for example if it prevents correct folding of ComP, which is required for PglS recognition and subsequent glycosylation (47), or partially blocks its access to the glycan attachment site.

de Alwis et al reported immunisation of rabbits with purified MdtA elicited a robust, protein-specific IgG response, which guided its selection as a carrier protein in this study (29). Glycosylation of novel carrier proteins risks masking of important protein epitopes with the glycan (24, 45). Bioconjugation offers an advantage over chemical conjugation for this process, as the site of glycosylation can be precisely determined by placement of the OST-recognition sequence. In our work, using C-terminal glycosylation, we confirm the immunogenicity of both glycosylated and unglycosylated MdtA in a mouse model, suggesting that both ComP fusion and glycosylation do not inhibit stimulation of an anti-MdtA IgG response. The mean IgG response against MdtAComP was higher compared to its glycosylated counterpart, although this difference was not significant. Notably, specific anti-ComP antibodies were also generated against both carriers, but this did not prevent the generation of a specific ExoA or MdtA response.

Selection and design of new glycoconjugate carrier proteins require multiple considerations such as immunogenicity, toxicity, manufacturability and regulatory approval. Additionally, as demonstrated here, it is necessary to characterise each novel carrier individually, as although the same C-terminal ComP tag was used for glycosylation of each protein, there was still variation in glycoconjugate yield and immune response. This could be further investigated through testing multiple configurations of each carrier protein, both in terms of number and position of glycosylation site(s). For example, glycoconjugate yield can be improved through inclusion of additional glycosylation sites, often achieved using multiple terminal “glycotags” (34). As was found in this study, this approach can be challenging when using large glycosylation tags such as the 114 amino acid ComP, but shorter ComP sequons have been identified that can still be glycosylated and could be more amenable to multiple copies (47).

Indeed, Knoot *et al.* recently demonstrated enhanced glycosylation of ExoA through incorporation of up to six, 23 amino acid ComP sites and demonstrated how position of these sites within the protein sequence can impact glycosylation (48). Furthermore, directed evolution of PglS generated a mutant capable of glycosylating two serine residues within a single ComP sequence, again providing opportunities for increased polysaccharide loading of carrier proteins (49).

Critically, both bioconjugates induced production of an anti-SSOAg IgG response, a key characteristic of effective *Shigella* glycoconjugate vaccines. A review of evidence obtained from both observational and vaccine efficacy studies found serum IgG antibodies against the *Shigella* LPS are associated with protection from Shigellosis (14). The anti-O-antigen response was approximately two-fold higher for SSOAg-ExoA compared to SSOAg-MdtA but the reason for this is unknown. Although both bioconjugates were highly glycosylated with minimal unglycosylated protein present, vaccine dose was based on protein concentration, therefore there may have been variation in the amount of glycan in each vaccine. Reglinski et al, investigated the use of novel carriers in the use of double-hit bioconjugate vaccines against *Streptococcus pneumoniae* serotype 4 and also found heterogeneity in the anti-capsular antibody response between the different carriers (25).

The detection of antibody binding to a chosen antigen is a key first step in the development of a novel vaccine candidate. The availability of suitable small animal models for use in *Shigella* challenge models has been limited but these are improving (50). Challenge experiments could aid in the understanding of protective efficacy of the bioconjugates vaccines developed here and indeed if this can be improved and/or provide heterologous *Shigella* protection through use of MdtAComP compared to ExoAComP as the glycan carrier. In summary, we demonstrate the heterologous production of a prototype, two-pronged bioconjugate vaccine against Shigellosis, by combining a conserved Shigella-specific protein and *S. sonnei* glycan antigen. This vaccine generated a Shigella-specific antibody response in a mouse immunisation model, thus demonstrating the flexibility and utility of bioconjugation for the generation of such vaccines.

## Acknowledgments

This work was supported by the Wellcome Trust (WT223838/Z/21/Z) and BactiVac. We thank Christian Harding and Mario Feldman for the constructs pCH4 and pACT3_pglS, Vanessa Terra and Mark Reglinski for their advice and discussions and Ian Passmore for critical reading of the manuscript.

