## supplementary material for "Development and evaluation of a dual target glycoconjugate vaccine against *Shigella sonnei*"

Supplemental Figures

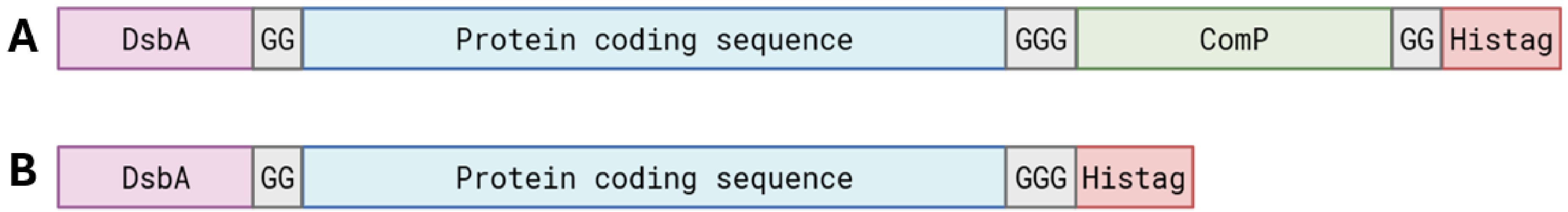


**Figure S1. Schematic of carrier protein design.** Each carrier protein (blue) was modified with a N-terminal DsbA signal sequence for trafficking to the periplasm (purple), double or triple glycine linkers (grey) and 6XHistag (red) for nickel-affinity purification. Carrier proteins varied by presence and number of ComP tags (green) for glycosylation with PglS. (A) C-terminal ComP tag and (B) no ComP tag.

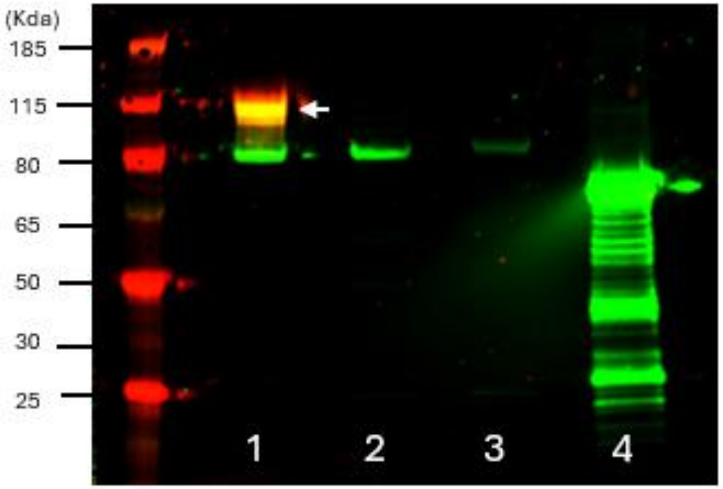


**Figure S2. Western blot analysis of PglS-mediated glycosylation of ExoAComP with the *S. sonnei* O-antigen in *E. coli* SDB1.** (1) *S. sonnei* O-antigen (SSOAg) expression plasmid (pBPSO) with ExoAComP and PglS, (2), no PglS control- pBPSO and ExoAComP only, (3) no glycan control- ExoAComP and PglS only, (4) pBPSO co-expressed with PglB oligosaccharyltransferase and compatible carrier protein, ExoA10. Nickel-affinity purified proteins were resolved by SDS-PAGE and Western Blot probed using anti-6xHistag antibody (green) and anti-SSOAg antibody (red/yellow). White arrow indicates glycoconjugate.

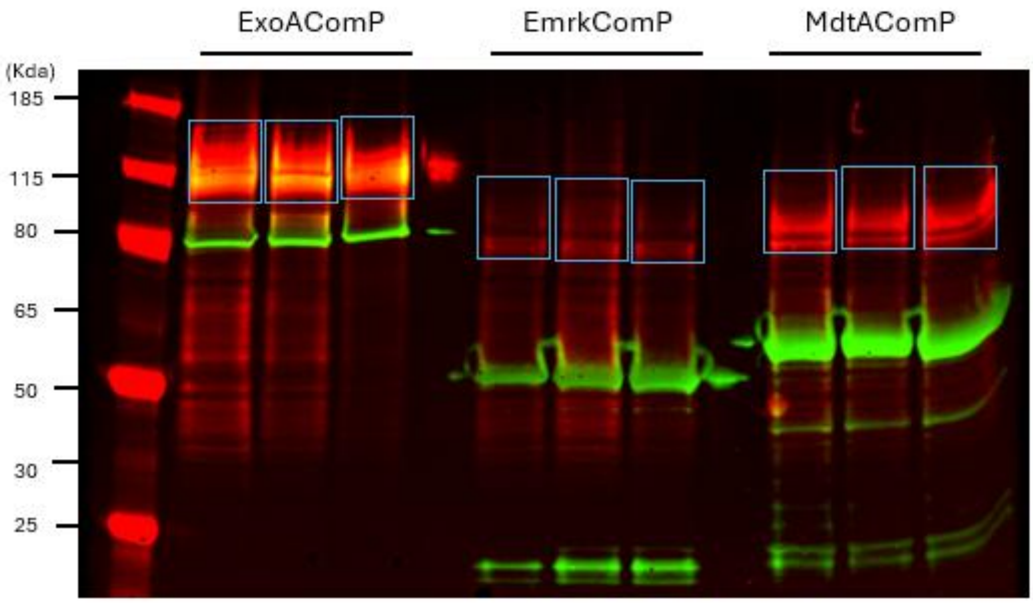


**Figure S3. Densitometry analysis of *Shigella sonnei* glycoconjugate production.** Three biological replicates of ExoAComP, EmrKComP and MdtAComP glycosylated with the *S. sonnei* O-antigen (SSOAg) were resolved by SDS-PAGE and detected via Western blot using anti-6xHistag antibody (green) and anti- SSOAg antibody (red). Blue boxes denote regions used for glycoconjugate quantification via densitometry using ImageJ software.

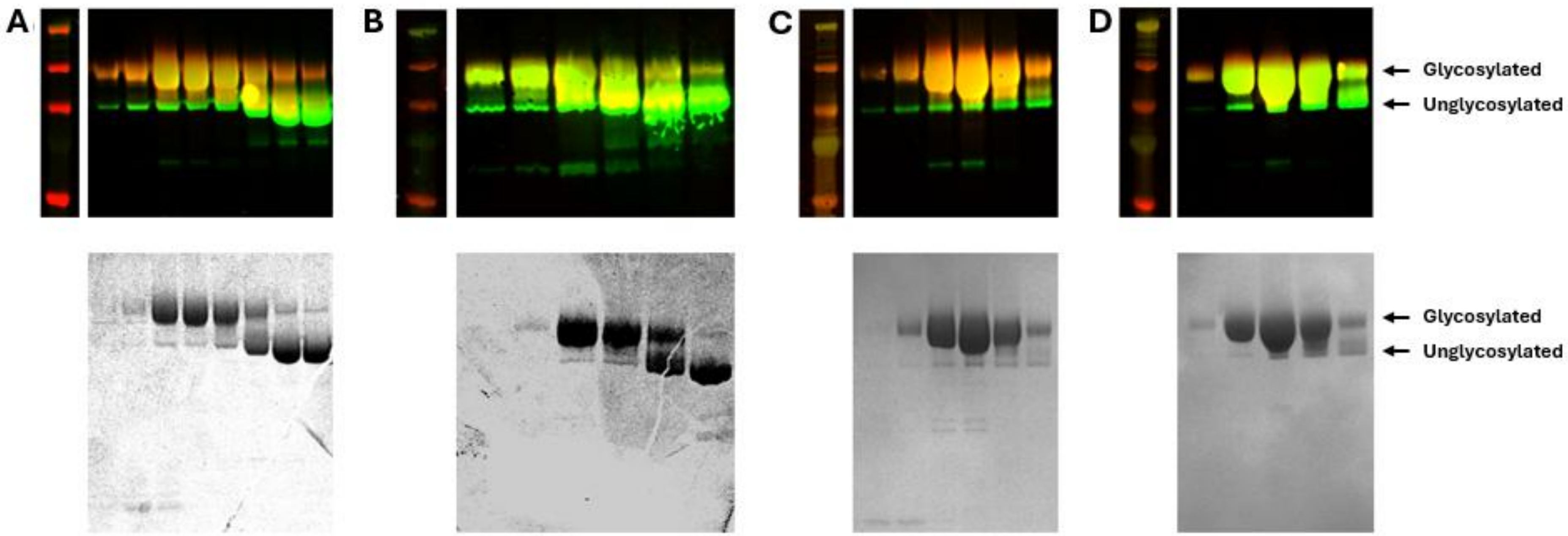


**Figure S4. Sequential anion exchange chromatography (AEX) for purification of *S. sonnei*-specific glycoconjugates.** Nickel-affinity purified ExoAComP conjugated to the *S. sonnei* O-antigen (SSOAg) was further purified via AEX for separation of glycosylated and unglycosylated protein using an increasing NaCl gradient. Elution fractions from each round of AEX were resolved by SDS-PAGE and visualised by Western blot (top panel) and Imperial staining (bottom panel). Fractions containing the most glycoconjugate were pooled and AEX repeated. Images A to D represent sequential rounds of AEX, from first to last, with each well containing a different elution fraction. Western blots were visualized using anti-6xHistag antibody (green) and anti- SSOAg antibody (red).

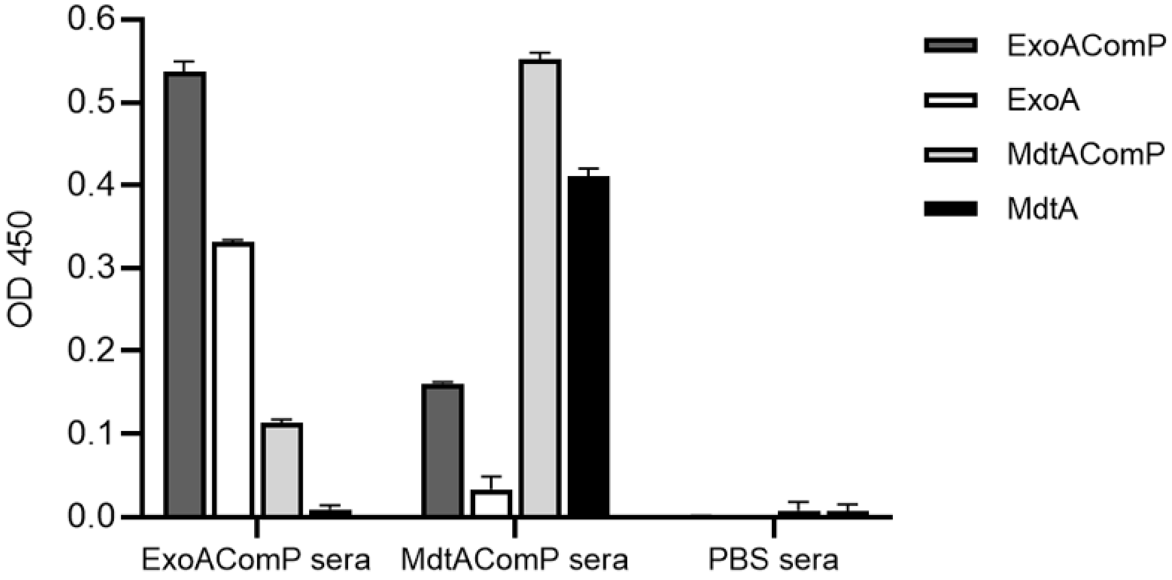


**Figure S5. Anti-carrier protein antibody response to ExoA and MdtA, with and without ComP.** ELISA quantification ($OD_{450}$) of IgG response to ExoA and MdtA, with and without ComP, in mice immunised with unglycosylated ExoAComP or MdtAComP, or PBS only. Each bar represents a different carrier protein, with pooled sera from each immunisation group on the x-axis. Error bars represent standard deviation from three technical replicates.

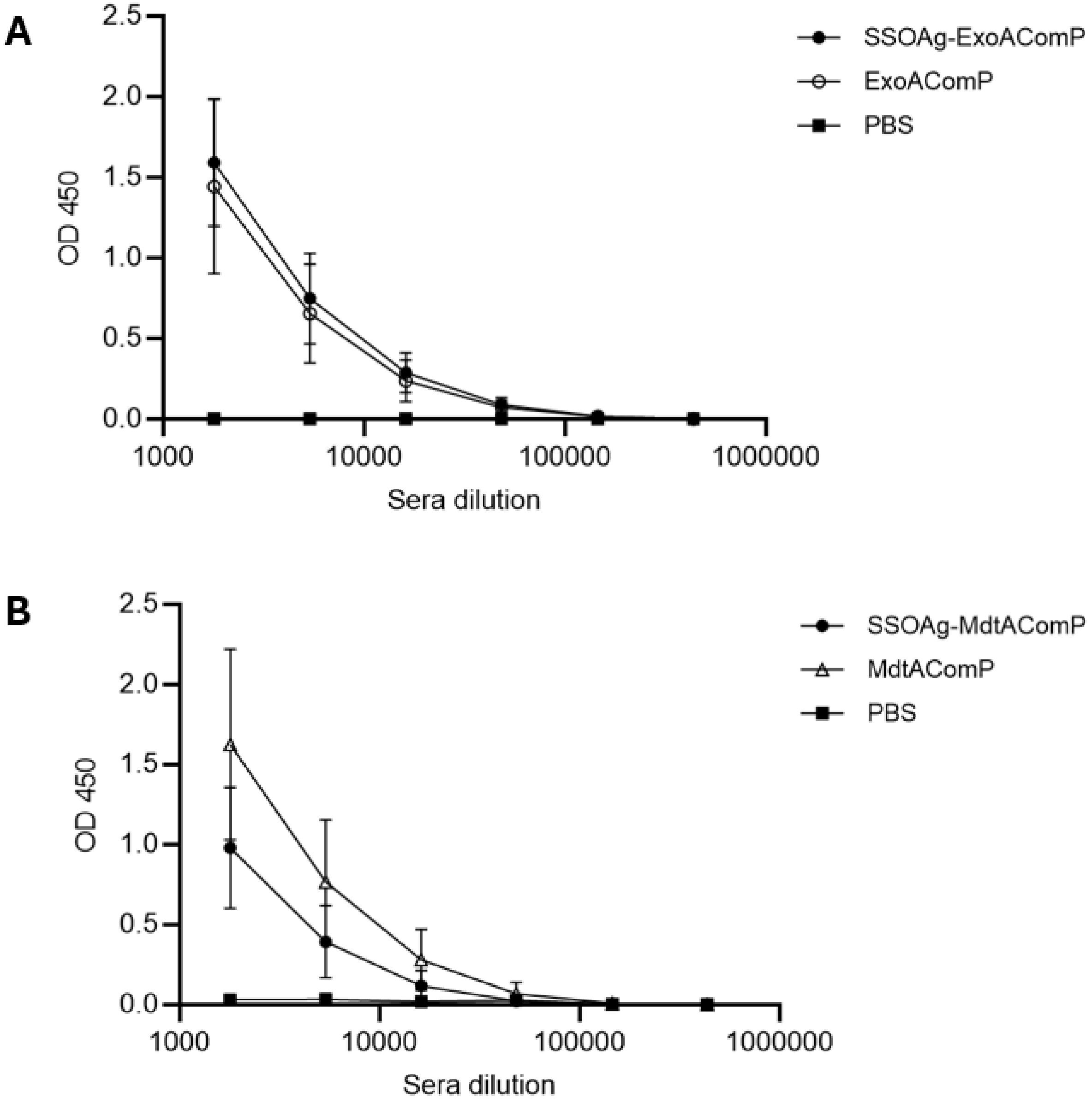


**Figure S6. Anti-carrier protein antibody response to ExoA and MdtA.** ELISA quantification of IgG response to ExoA and MdtA in mice immunised with glycosylated or unglycosylated (A) ExoAComP or (B) MdtAComP. $OD_{450}$ is reported for 6-point, 3-fold dilution series of sera, error bars represent standard deviation for seven biological replicates.

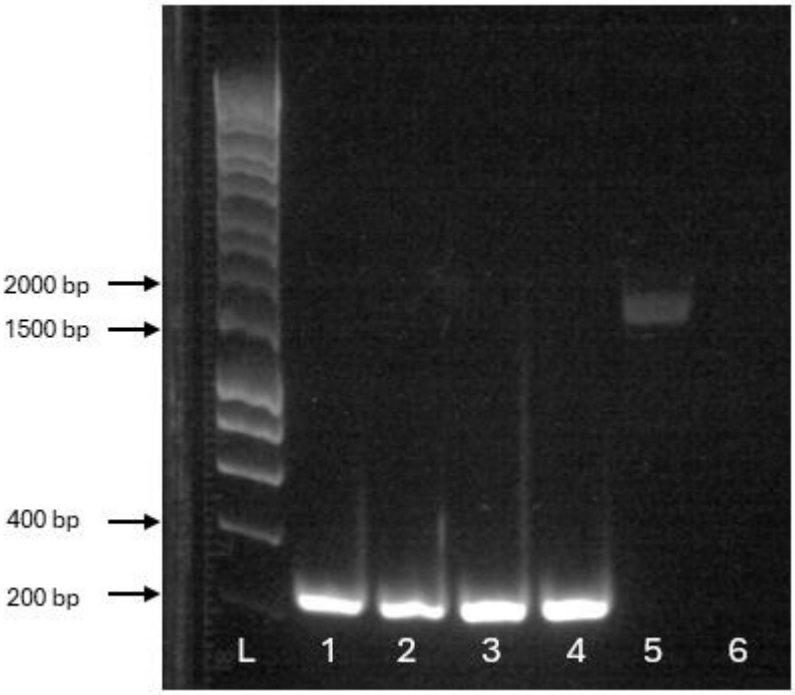


**Figure S7. Generation of W3110Δ*mdtA*.** *mdtA* was deleted from the W3110 chromosome using lamda red recombineering. *mdtA* was replaced with a chloramphenicol cassette which was flanked by *dif* recombinase sites for excision by the *E. coli* Xer recombinase. PCR using primers flanking the *mdtA* gene were used to screen colonies which had lost chloramphenicol resistance and amplicon resolved by gel electrophoresis. (1-4) W3110 Δ*mdtA* clones, (5) W3110 wild-type, (6) d$H_2O$ negative control. Wild-type W3110, 1529 bp, W3110Δ*mdtA,* 309 bp.

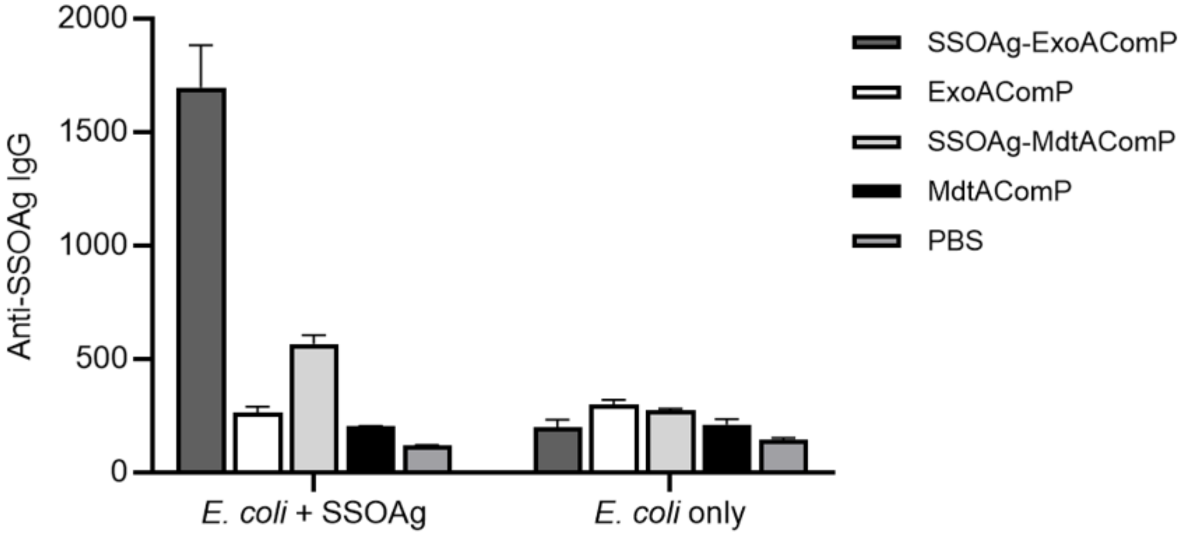


**Figure S8.** Antibody response to *S. sonnei* O-antigen. ELISA quantification of IgG response to whole *E. coli* cells expressing *S. sonnei* O-antigen or empty vector (no O-antigen) in mice immunised with glycosylated or unglycosylated ExoAComP or MdtAComP. A 6-point, 3-fold dilution series was tested for pooled serum from each group and IgG response reported as the reciprocal serum dilution that gave an OD 0.2. Error bars represent standard deviation for three technical replicates. SSOAg, *S. sonnei* O antigen.

Supplemental Tables

**Table S1. List of strains used in this study.**

| **Strain** | **Genotype** | **Source** |
| --- | --- | --- |
| *E. coli* W3110 | F- *mcrA mcrB* In(*rrnD-rrnE*)1 | ATCC 27325 |
| *E. coli* W3110Δ*mdtA* | | This study |
| *E. coli* SDB1 | W3110 Δ*waaL*, Δ*wecA* | (1) |
| *Plesiomonas shigelloides* O17 | O17 O-antigen | NCTC10360 |
| *E. coli* EC100D (*pir*+) | *$F^-$ mcrA Δ(mrr-hsdRMS-mcrBC) ϕ80dlacZΔM15 ΔlacX74 recA1 endA1 araD139 Δ(ara, leu)7697 galU galK λ- rpsL ($Str^R$) nupG $pir^+$(DHFR)* | (2) |

**Table S2. List of plasmids used in this study.**

| **Plasmid** | **Description** | **Source** |
| --- | --- | --- |
| pBBR1MCS-3 | Cloning vector ($Tet^R$) | (3) |
| pBPSO | pBBR1MCS-3 containing 12.2 Kb *Plesiomonas shigelloides* O17 O-antigen region from *wzz* to *wbgZ*, inserted into ApaI site. | This study |
| pACT3_pglS | pACT3 ($Cm^R$) containing *Acinetobacter baylyi pglS* | (4) |
| pEXT22_pglS | pEXT22 ($Kan^R$) containing *Acinetobacter baylyi pglS,* inserted into EcoRI and BamHI sites. | This study |
| pEC415 | Expression vector ($Amp^R$), L-arabinose inducible | (5) |
| pEC415_ExoAComP | pEC415 containing detoxified *Pseudomonas aeruginosa* ExoA with N-terminal DsbA periplasmic signal sequence and C-terminal ComP sequon and 6xHistag. | This study |

| | | |
|---|---|---|
| pEC415_EmrKComP | pEC415 containing EmrK (amino acids 38-387) with N-terminal DsbA periplasmic signal sequence and C-terminal ComP sequon and 6xHistag. | This study |
| pEC415_MdtAComP | pEC415 containing MdtA (amino acids 66-453) with N-terminal DsbA periplasmic signal sequence and C-terminal ComP sequon and 6xHistag. | This study |
| pCH4 | pEXT20 containing detoxified *Pseudomonas aeruginosa* ExoA with N-terminal DsbA periplasmic signal sequence and C-terminal ComP sequon and 6xHistag. | (6) |
| pEXT20 | Expression vector ($Amp^R$), IPTG-inducible | (7) |
| pEXT20_ExoA | pEXT20 containing detoxified *Pseudomonas aeruginosa* ExoA with N-terminal DsbA periplasmic signal sequence and 6xHistag. | This study |
| pEXT20_MdtA | pEXT20 containing MdtA (amino acids 66-453) with N-terminal DsbA periplasmic signal sequence and 6xHistag. | This study |
| pKD46 | Cloning vector ($Amp^R$), containing lamda red recombineering machinery, temperature sensitive. | (8) |
| pKD4 | Cloning vector ($Amp^R$,$Kan^R$) used for construction of *mdtA* mutagenesis cassette. | (8) |
| pKD4_mdtA | pKD4 containing *mdtA* mutagenesis cassette. | This study |

**Table S3. List of oligonucleotides used in this study.**

| Oligonucleotide | Description | Sequence |
|---|---|---|
| PsOFApaI | Amplification of 12.2Kb region of *P. shigelloides* O-antigen locus, from *wzz* to *wbgZ* | tatcatGGGCCCgcagttggcgatatcctgtt |
| PsORApaI | | aatcatGGGCCCtcacaggcgtatgacattgc |
| pEC415DsbA_f | Amplification of pEC415 for insertion of *exoAComP*, *emrKComP*, and *mdtAComP* expression constructs | attgagaattcttgaagacgaaa |
| pEC415DsbA_r | | ttcattatgttattcctccttattta |
| ExoA_NComPdel_f | Removal of N-terminal ComP sequon from pEC415_ExoAComP | gcggccgaggaggcattcgatttatggaatgaatgtgc |
| ExoA_NComPdel_r | | tccgccctgcgccgcgct |
| EmrK_NComPdel_f | Removal of N-terminal ComP sequon from pEC415_EmrKComP | ggcggtgaactggaagacatgattagtacggat |
| EmrK_NComPdel_r | | ctgcgccgcgctagcgctaaa |
| MdtA_NComPdel_f | Removal of N-terminal ComP sequon from pEC415_MdtAComP | ggcggtggccgtaacgactca |
| MdtA_NComPdel_r | | ctgcgccgcgctagcgctaaa |
| pEXT20_noComP_f | Amplification of pEXT20 for insertion of *exoA* and *mdtA* without ComP | caagcttctgttttggcgg |
| pEXT20_noComP_r | | gaattctgtttcctgtgtgaaatg |
| pKD4MdtA_f | Amplification of pKD4 for insertion of *mdtA* mutagenesis cassette | atatggaccatggctaattcccatg |
| pKD4MdtA_r | | gcatgcaagcttggcactg |
| MdtAcassette_f | Amplification of *mdtA* mutagenesis cassette from pKD4_mdtA | atgtgcccgtcattcagacg |
| MdtAcassette_r | | ggcgcgataaccgataatccc |
| MdtAscreen_f | Screening to confirm *mdtA* deletion from *E. coli* W3110 chromosome | ctcgcaaatgtgcccgtcatt |
| MdtAscreen_r | | ggatagtccacttccggcag |

Capital letters indicate restriction sites.
